# Decoding mesolimbic dopamine transmission in the olfactory tubercle and its contribution to methamphetamine responses through neurochemical sensing and chemogenetics

**DOI:** 10.1101/2024.11.12.622180

**Authors:** Rohan V. Bhimani, Ryan C. Pauly, Caroline E. Bass, Jinwoo Park

## Abstract

Central dopamine (DA) innervation of the olfactory tubercle (OT) from the ventral tegmental area (VTA) plays a critical role in encoding multisensory information and generating behavioral outputs necessary for survival. However, due to anatomical restrictions and the neurochemical heterogeneity of the VTA and OT, very little is known about the functional link between mesolimbic VTA-DA transmission in the OT and its role in mediating reward and drug seeking. In this study, we integrated *in vivo* fast-scan cyclic voltammetry with chemogenetics to (1) characterize the effects of chemogenetic modulation (excitation and inhibition) of mesolimbic DA transmission in the OT of both anesthetized and awake-behaving wild-type rats and (2) demonstrate that inhibition of VTA-DA neurons is sufficient to suppress methamphetamine-induced DA transmission as well as its locomotor and rewarding effects. These results offer novel insights into mesolimbic DA transmission in the OT and its contribution to substance use disorders.

---

The mesolimbic dopamine (DA) system consists of DA neurons in the ventral tegmental area (VTA) that project diffusely throughout limbic forebrain areas, including the ventral striatum, and plays a major role in locomotion, learning, and substance use^1, 2^. This system is also critically involved in diverse behavioral and physiological processes while its dysfunction is associated with many mental disorders and neurodegenerative diseases^3–6^. Preclinical work has shown that the major components of ventral striatum, the nucleus accumbens (NAc) and olfactory tubercle (OT), receive the densest VTA-DA innervation in the brain^7–9^. VTA-DA signaling in the NAc has been extensively studied for its roles in reward, motivation, and drug addiction^2, 10, 11^. In contrast, the OT, which is located ventral to the NAc, has often been overshadowed and considered simply an extension of the NAc rather than its own individual structure. However, accumulating studies have highlighted that the OT is a distinct part of the ventral striatum and VTA-DA innervation of the OT (specifically the medial subregion) plays a key role in encoding reward and may be more susceptible to the effects of psychostimulants than the NAc^7, 8, 12–14^. For example, studies have shown that systemic administration of methamphetamine (METH), a monoamine transporter inhibitor and the second most illicitly used amphetamine-type stimulant, induces greater c-Fos reactivity in the OT of a rat than any other limbic brain area^15^ and rats self-administer cocaine more readily into the medial OT than other striatal subregions^16^.

Despite this evidence, very little is known about the functional roles of VTA-DA transmission in the OT and its contributions to reward and drug seeking due to technical limitations. Structurally, the rat OT is a ∼100-400 µm trilaminar region located at the ventral most portion of the striatum and is adjacent to several other DA-rich brain areas including NAc and caudate putamen^8, 13^. The OT itself can be subdivided into lateral and medial regions each with distinct patterns of DA signaling, with the medial(m) OT considered to play a greater role in reward^13, 17^. Traditional methods to study brain regions such as chemical lesion, pharmacological manipulations, and electrical stimulation are often non-selective, off-target, or have non-specific effects^18–20^, making it very challenging to selectively modulate VTA-DA neurons and their projections exclusively in the neurochemically heterogeneous VTA^21, 22^. Determining the distinct roles of DA transmission within such discrete brain areas requires approaches that allow for precise control of VTA-DA neurons as well as the ability to selectively monitor DA transmission at their terminals such as the subregions of the OT.

In this study we integrated cell-type specific chemogenetics using Designer Receptors Exclusively Activated by Designer Drugs (DREADDs) with real-time neurochemical sensing via fast-scan cyclic voltammetry to modulate VTA-DA signaling and monitor its transmission in the medial OT (mOT), respectively. These DREADDs are activated by the relatively pharmacologically inert ligand Clozapine-N-oxide (CNO), however, it is not well understood until now how chemogenetic modulation of VTA-DA neurons impacts terminal DA transmission *in vivo*. Herein, we (1) determined how DREADDs modulate extracellular VTA-DA regulation (release and clearance) in the mOT of anesthetized rats after optimizing chemogenetic parameters such as CNO dose and drug time course, (2) identified the effects of chemogenetic modulation of VTA-DA neuronal activity on DA transmission in the mOT of awake behaving rats, and (3) determined the role of VTA-DA neurons in METH-induced DA transmission in the mOT. Through the integration of neuroanatomical, behavioral, neurochemical, and pharmacological approaches, our results demonstrate for the first time (1) a dose response for CNO’s ability to modulate VTA-DA neurons expressing DREADDs and DA transmission, (2) how chemogenetic manipulation of VTA-DA neurons affects DA transmission, and (3) inhibition of VTA-DA neuronal activity can attenuate METH-induced DA transmission in the mOT as well as reduce the locomotor and rewarding properties of METH. Importantly, all experiments were conducted and validated in “*wild-type”* rats. These results highlight a correlation between VTA-DA transmission in the mOT and METH seeking and sets the guidelines for the proper use of chemogenetics to decode the distinct roles of mesolimbic DA across striatal subregions.

## Results

### DREADD expression in VTA-DA neurons and its terminals in the OT

A schematic of the combinatorial viral targeting system used to selectively transduce catecholamine neurons in wild-type rats is shown in Figure 1A^23^. DREADD expression was primarily restricted to VTA-DA neurons (Figure 1B & C) and demonstrated robust co-expression with tyrosine hydroxylase (TH) throughout terminal regions in the NAc and OT (Figure 1D & E) five weeks after virus infusion. Over 99.6 % of mCherry+ cells co-expressed with TH in the VTA as verified by confocal microscopy (*Mann Whitney*, *p =* 0.0143, n = 4 rats). The boundary between the NAc and OT is clearly indicated by a lack of TH expression in Figure 1D and Supplementary Figure 1. Expression of both TH and mCherry are observed throughout the medial and lateral OT. It is important to note that similar to the NAc, over 94 % of the catecholamine content in the OT is DA at this coordinate^8, 9, 13, 24^.

**Figure 1:**
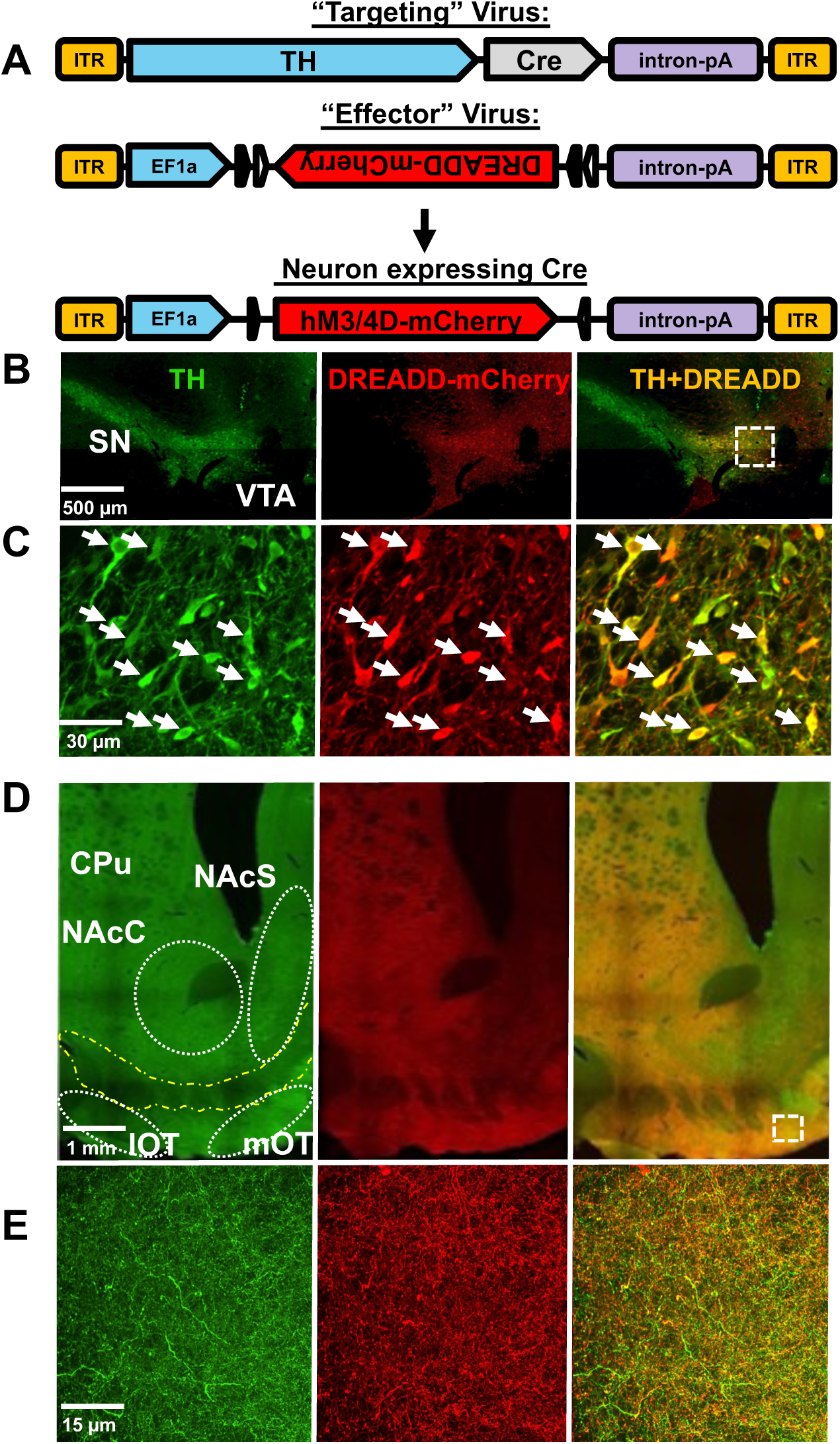
**A.** Schematic of the combinatorial viral targeting system. **B-E**. DREADD expression in VTA-DA cell bodies and their terminals. Immunofluorescence of TH (green) and DREADD-mCherry (red) expression in the VTA (**B-C**) and ventral striatum (**D-E**) demonstrating robust co-expression (orange). White dotted squares in Fig. B and D indicate area of magnification in Fig. C and E, respectively. White arrows indicate mCherry+ neurons that co-express TH in the VTA. Strong overlap of TH+ terminals and RFP is observed in the OT (D & E, right panel) CPu: Caudate Putamen; NAcC/S: Nucleus Accumbens Core/Shell; m/lOT: Medial/Lateral Olfactory Tubercle; SN: Substantia Nigra; VTA: Ventral Tegmental Area

### Temporal and dose-dependent effects of CNO on chemogenetic modulation of VTA-DA transmission in the OT of anesthetized rats

Very little is known regarding how chemogenetic modulation of VTA-DA neurons impacts terminal extracellular DA transmission *in vivo*^25^. In the following experiments, rapid changes in VTA-DA transmission at carbon-fiber microelectrodes (neurochemical sensors) coupled with fast-scan cyclic voltammetry (FSCV) were monitored in urethane-anesthetized rats 4-5 weeks after viruses were infused into the VTA of DREADD and mCherry control rats (Figures 2 & 3). This anesthetized approach is widely used to accurately characterize the effects of pharmacological agents on DA regulation (release and clearance) without confounding issues present in awake behaving rats during recording sessions^12, 26, 27^. In these experiments, a neurochemical sensor was lowered into the mOT until reproducible DA release evoked by electrical stimulation (20 Hz, 60 pulses) of the VTA was detected. A neurochemical and anatomical boundary exists between the NAc and OT where DA release is not observed (Figure 1 & Supplementary Figure 1), which allows us to determine DA release in the mOT reliably, as demonstrated in our previous studies^9, 13^. Once both stimulating and neurochemical sensing electrode placements were optimized in the VTA (8.5 – 9.0 mm ventral from the skull) and mOT (8.2 – 8.5 mm ventral from the skull), respectively, they were not changed for the duration of the experiment. Electrically evoked maximal DA concentration ([DA]_max_) and its clearance time (t_1/2_, time for [DA]_max_ to decay to half of its maximum) in the mOT were measured every 10 minutes before and after systemic administration of vehicle or CNO. Cumulative doses of CNO (consisting of 0.3 mg/kg IP administered followed an hour later by a second dose of either 1.0 or 3.0 mg/kg IP) were given to anesthetized rats expressing mCherry (no DREADD control), hM3D (excitatory), or hM4D (inhibitory). These CNO doses (0.3, 1.0, and 3.0 mg/kg, IP) are based on those used in previous studies examining DA-related behaviors in rats and mice expressing DREADDs in VTA-DA neurons^28, 29^. Figure 2A shows representative color plots and concentration vs time traces for electrically evoked DA after vehicle (1.5 % ethanol in saline, 1.2 mL/kg) and a pair of cumulative doses of CNO (0.3 and 1.0 mg/kg, IP) in mCherry (top), hM3D (middle) and hM4D (bottom) rats. The voltammogram in the inset identifies the signal as catecholamine. CNO effects on electrically evoked DA release were shown to maximize after 30-40 minutes in both hM3D and hM4D groups (Supplementary Figure 2). CNO did not cause any significant change in electrically evoked DA regulation in mCherry rats at any dose (0.3 mg – 3.0 mg, Figure 2 & Supplementary Figure 3). In hM3D rats, CNO dose-dependently enhanced [DA]_max_ (*F*(3, 20) = 58.35, *p* < 0.0001) and t_1/2_ (*F*(3, 22) = 9.179, *p* = 0.0004). 0.3 mg/kg CNO caused a significant increase in both electrically evoked [DA]_max_ (86.1 ± 10.7 %) and its t_1/2_ (33.9 ± 8.4 %) whereas 1.0 mg/kg further enhanced DA signaling ([DA]_max_ = 164.7 ± 14.7 %, t_1/2_ = 58.6 ± 16.2 %) (Figure 2B). There was no significant difference in [DA]_max_ and t_1/2_ between 1.0 and 3.0 mg/kg CNO (Figure 2B and Supplementary Figure 3). In contrast, [DA]_max_ significantly decreased at all doses of CNO in hM4D rats (*F*(3, 17) = 49.53, *p* <0.0001) without effecting the t_1/2_ (*F*(3,27) = 0.9717, *p* = 0.4205). 0.3 mg/kg CNO showed maximized suppression of DA release ([DA]_max_= -64.6 ± 3.5 %) without further inhibition at higher CNO doses (1.0 and 3.0 mg/kg). Interestingly, 1 hour after CNO administration, the rate of rise (slope indicated in blue line) of DA concentration during electrical stimulation significantly increased in hM3D rats (Vehicle = 381 ± 6.9 nM/s; CNO 0.3 mg/kg = 548 ± 28 nM/s; CNO 1.0 mg/kg = 637± 4.9 nM/s, *F*(2,9) = 6.633, *p* = 0.017) (Figure 2A and Supplementary Figure 3). In contrast, the rate of rise of DA concentration significantly decreased in hM4D rats after CNO (Vehicle = 398 ± 105 nM/s and 0.3 mg CNO = 28.3 ± 2.5 nM/s, *t*(6)=4.331, *p* = 0.0025). The decreased slopes of DA concentration were similar across all doses of CNO in inhibitory rats. Additionally, the latency (time for the onset of DA release from electrical stimulation) increased after CNO (0.4 ± 0.09s, n = 4 rats) in hM4D rats, whereas mCherry and hM3D rats did not show a delay in DA release following stimulation.

**Figure 2:**
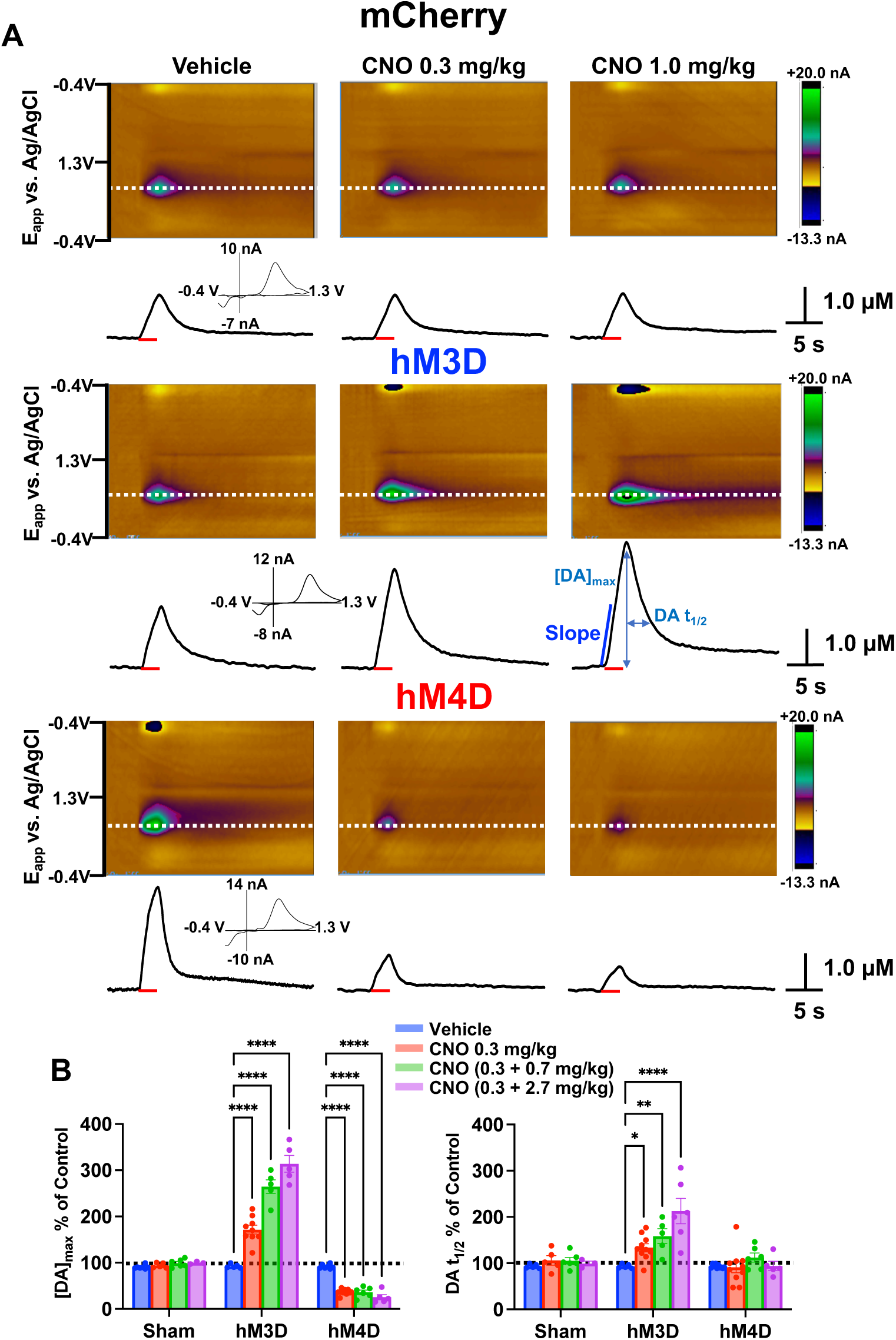
**A.** Representative color plots (top) and dopamine (DA) concentration vs time traces at the oxidation potential for DA (dotted white line) in the OT of anesthetized mCherry and DREADD rats following electrical stimulation (20 Hz, 60 pulse; denoted by red bar) of the VTA before and after CNO administration. Voltammograms in the inset identify the signal as catecholamine **B.** Average changes in maximally evoked DA concentration ([DA]_max_, left) and the time for [DA]_max_ to decay to half of the maximum (t_1/2_, right) following different doses of CNO. *p < 0.05,**p < 0.01,***p < 0.001, ****p < 0.0001.

**Figure 3:**
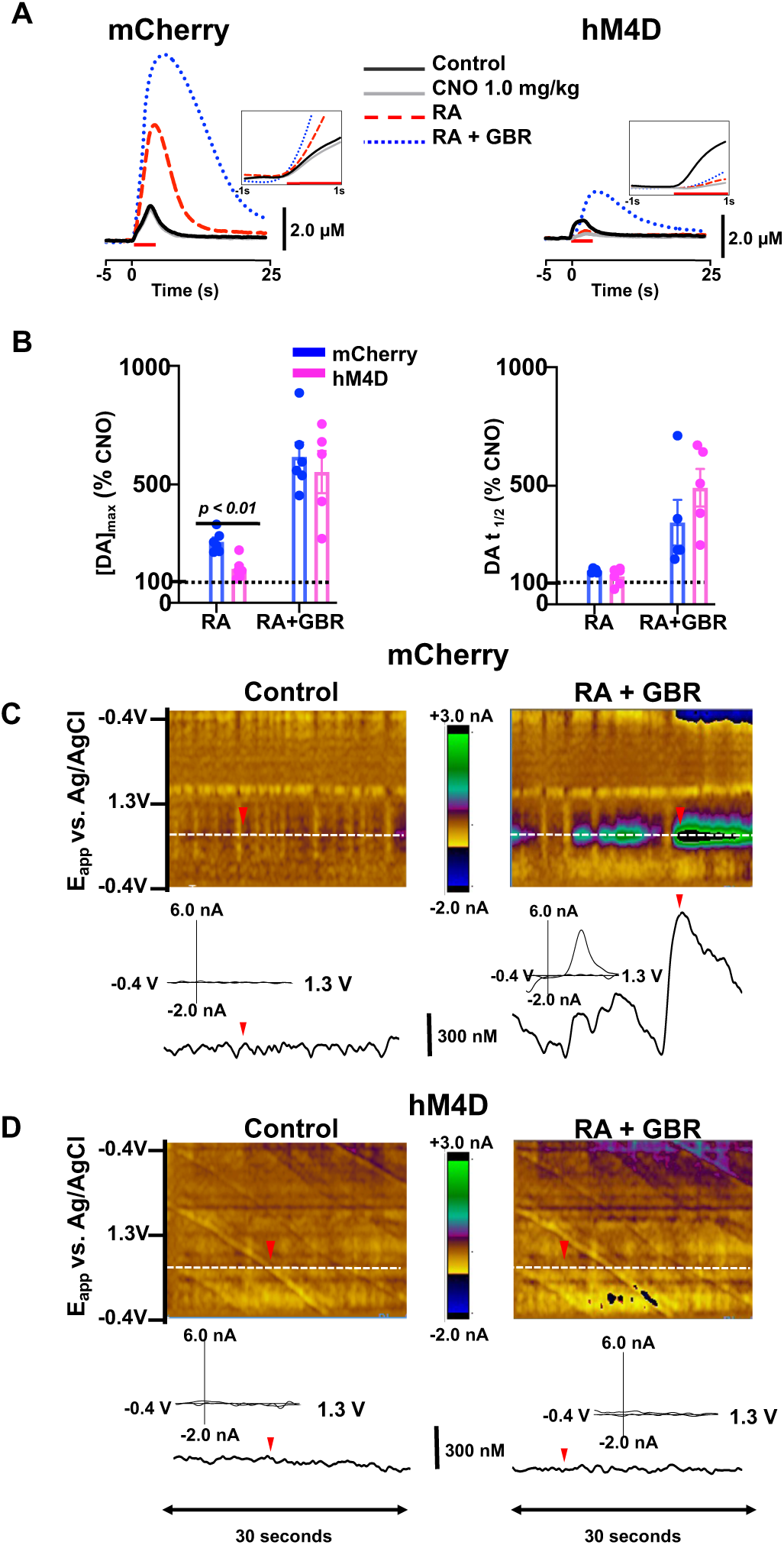
**A.** Representative dopamine (DA) concentration vs time traces in anesthetized mCherry (left) and hM4D (right) rats following systemic administration of CNO (1.0 mg/kg), raclopride (RA, 2.0 mg/kg) and GBR 12909 (GBR, 15.0 mg/kg). Inset shows longer latency to release DA following CNO administration in hM4D but not mCherry rats. **B.** Average maximal change in electrically evoked [DA]_max_ (left) and t_1/2_ (right) following RA, and RA+GBR. Comparisons made against CNO as baseline. Electrical stimulation (20 Hz, 60 pulses) of the VTA is denoted by red bar. Representative color plots (top) and concentration vs time traces (bottom) at the oxidation potential (∼0.67 V) for DA (dotted white line) in mCherry (**C**) and hM4D (**D**) rats before drugs and after co-administration of CNO (1.0 mg/kg), raclopride (RA, 2.0 mg/kg) and GBR 12909 (GBR, 15.0 mg/kg). Inverted red triangles indicate the time when a background subtracted voltammogram was extracted from to identify the analyte.

### Effects of pharmacological manipulations on chemogenetically inhibited DA regulation

After characterizing how chemogenetic inhibition of VTA-DA impacts electrically evoked DA transmission, its inhibitory effects on DA regulatory mechanisms (release and clearance) in the mOT of urethane-anesthetized rats were determined using well-characterized pharmacological manipulations of DA D2 autoreceptors and transporters^13, 30^. Figure 3A shows representative electrically evoked DA concentration vs time traces in the mOT of mCherry and hM4D rats that were systemically administered CNO (1.0 mg/kg) followed by the D2 autoreceptor antagonist, raclopride (RA, 2.0 mg/kg) and DA transporter inhibitor (DAT), GBR 12909 (GBR, 15.0 mg/kg). In mCherry rats, the D2 autoreceptor antagonist increased electrically evoked [DA]_max_ and the DAT inhibitor further significantly potentiated both [DA]_max_ and DA t_1/2_ (*t*(10) = 5.632, *p* < 0.0001). In contrast, hM4D rats showed no significant increase in [DA]_max_ after raclopride administration, however, GBR caused a significant increase in [DA]_max_ (*t*(10) = 3.217, *p* = 0.0046) compared to raclopride alone. There was no significant difference in the GBR effects on both [DA]_max_ and t_1/2_ in mCherry and hM4D rats (Figure 3B). Magnified DA concentration traces one second before and during the first second of electrical stimulation are shown in the inset in Figure 3A. Antagonism of the D2 receptor did not reverse the CNO-induced increased latency, the (delay of DA release, 0.3 ± 0.05 s) or decreased slope (44.5 ± 23 nM/s) in inhibitory rats. Following administration of the DA transporter inhibitor, the rate of rise of DA concentration was significantly higher in mCherry rats (978 ± 306 nM/s) compared to hM4D rats (567 ± 85 nM/s) (*t*(6) = 8.012, *p* = 0.0002). However, the longer latency (0.4 ± 0.09s, n = 4 rats) for release during electrical stimulation induced by CNO was not reversed after DAT blockade (0.18 ± 0.17 s) in hM4D rats (Figure 3A).

Naturally occurring phasic DA transients that are implicated in reward, learning, locomotion, and underlie many goal-directed behaviors occur in awake-behaving rats^31–33^. Although rarely observed in anesthetized rats, spontaneous DA transients in the mOT have been clearly monitored after co-administration of DA autoreceptor and transporter inhibitors^13, 34, 35^. mCherry rats showed robust phasic DA transmission after RA+GBR following CNO (1.0 mg/kg) similar to our previous studies in naïve rats^9, 35^(Figure 3C). In contrast, hM4D rats did not exhibit any detectable transients (S/N ≥ 5) following blockade of the DA autoreceptors and transporters (n=4) (Figure 3D). The voltammograms in the inset identify the signals as catecholamine in origin (denoted by red triangles).

### Effects of chemogenetic modulation of VTA-DA neurons on locomotion and phasic DA transmission in the mOT

We next determined how chemogenetic excitation/inhibition of VTA-DA in awake-behaving rats modulates locomotion by different doses of CNO. Vehicle (1.2 mL/kg) followed by cumulative doses of CNO (0.3, 1.0, and 3.0 mg/kg) were administered 40 minutes apart. This time interval was selected based on previous behavioral studies^28, 36^ and our neurochemical data (Figure 2 and Supplementary Figure 2 & 3) showing that CNO effects are maximized and do not significantly change any further ∼ 40 min after systemic administration. Figure 4A shows representative horizontal locomotor tracks after vehicle (1.2 mL/kg, left) and CNO 1.0 mg/kg (right). Following cumulative doses of CNO, hM3D rats demonstrated a significant, dose-dependent increase in locomotion (*F*(3, 25) = 7.076, *p* = 0.0013) and rearing (an index of exploratory activity, *F*(3, 25) = 8.05, *p* = 0.0017)(Figure 4B). 1.0 and 3.0 mg/kg of CNO dose responsively increased the total distance traveled compared to vehicle and 0.3 mg/kg CNO, while 0.3, 1.0 and 3.0 mg/kg CNO all increased vertical counts. However, there was no significant difference between the higher doses. In contrast, hM4D rats demonstrated a slight decrease in locomotion and rearing, but it was not significantly different from mCherry rats, which did not show any changes in horizontal locomotion or rearing at after CNO.

**Figure 4:**
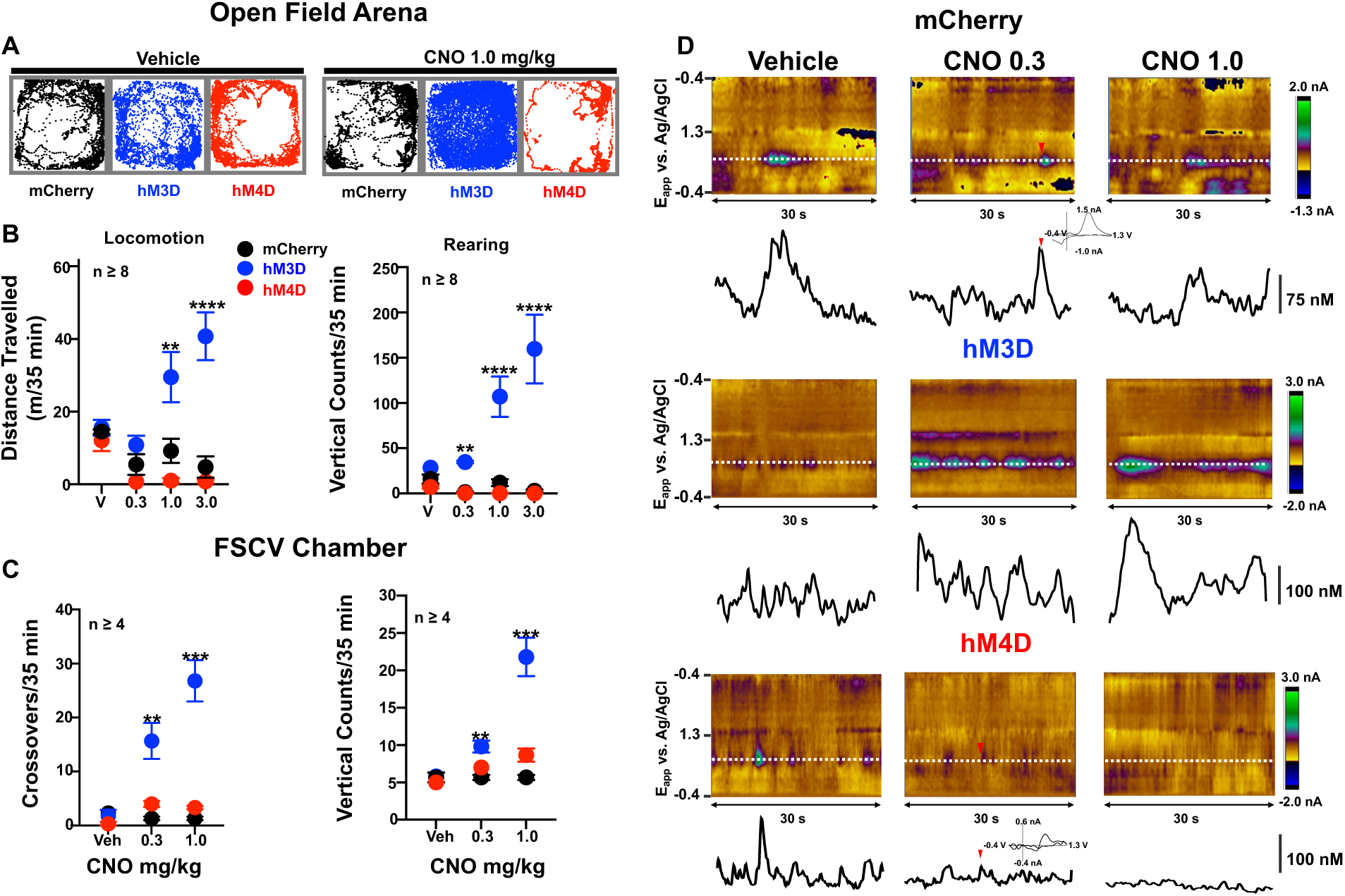
**A.** Representative tracks of total distance travelled in mCherry, excitatory (hM3D), and inhibitory (hM4D) rats after systemic administration of Vehicle (left) and CNO (1.0 mg/kg, right). Average changes in horizontal locomotion (left) and rearing (right) activity monitored every 40 minutes following cumulative doses of CNO in the open field arena (**B**) and FSCV chamber (**C**). **D.** Representative concentration vs time traces at the oxidation potential for DA (dotted white line, color plot) in mCherry, hM3D, and hM4D rats after vehicle (1.2 mL/kg) and CNO (0.3 and 1.0 mg/kg). Inverted red triangles denote phasic DA transients where cyclic voltammograms are extracted. Inset: cyclic voltammogram for catecholamine. **p < 0.01,***p < 0.001, ****p < 0.0001.

To the best of our knowledge, it has not been determined until now how chemogenetic excitation/inhibition of VTA-DA neurons modulate naturally occurring DA transients in its terminal regions, including the mOT, and the associated behavioral outputs. In this study, the effects of chemogenetic modulation of VTA-DA neurons on phasic DA transmission in the mOT and behavior were investigated in custom-built FSCV recording chambers that were modified to resemble an open field arena. Behaviors were videotaped and scored manually following experiments. A neurochemical sensor was lowered in 200-300 µm increments from the medioventral NAc (DV – 6.0 mm) to the mOT (DV -8.0 ∼ 8.5 mm) by monitoring DA release evoked by electrical stimulation of the VTA to properly position the sensor in the mOT. mCherry, hM3D, and hM4D rats received a pair of cumulative doses (0.3 & 1.0 mg/kg, IP) of CNO (40 minutes apart). The two doses were selected for these experiments based on the anesthetized FSCV and behavior data showing no significant differences between 1.0 and 3.0 mg/kg CNO on electrically evoked DA in the OT and locomotion, respectively (Figure 2 & 4B). Representative color plots and DA concentration vs. time traces in mCherry (top), hM3D (middle) and hM4D (bottom) rats ∼40 minutes after vehicle and CNO are shown in Figure 4D. Importantly, at both doses, CNO did not impact naturally occurring phasic DA transmission in mCherry rats. The voltammogram in the inset from a single timepoint (indicated by red triangles) identifies the signal as catecholamine.

hM3D rats showed an increase in DA transient intensity (concentration, nM) following CNO administration (Table 1, *F*(2,8) = 22.63, *p* = 0.0005). Transient frequency (Hz) also significantly increased with 0.3 mg/kg CNO but was not enhanced more with 1.0 mg/kg (*F*(2,8) = 7.56, *p* = 0.0143). At 1.0 mg/kg, there were often overlapping transient peaks that prevented accurate analysis of DA transient frequency (Figure 4D, middle). In contrast, for hM4D rats, DA transient intensity was significantly inhibited after CNO (0.3 mg/kg) and further decreased with 1.0 mg/kg (*F*(2,8) = 22.63, p = 0.0005) (Figure 4D, Table 1). The transient frequency also significantly decreased at 0.3 and 1.0 mg/kg (*F*(2,6) = 118.3, *p* < 0.0001). Like in the open field chambers (Figure 4B), locomotion and rearing in the custom built FSCV chambers for hM3D rats also increased dose dependently but there were no significant changes in the hM4D rats, though we could not rule out a floor effect for some metrics (Figure 4B).

**Table 1:** Average change in dopamine (DA) transient concentration (nM) and frequency (Hz) in DREADD rats following cumulative administration of vehicle or CNO 40 minutes apart.

| DREADD |  | Vehicle | CNO 0.3 mg/kg | CNO 1.0 mg/kg |
| --- | --- | --- | --- | --- |
| mCherry | Concentration (nM) | 44.5 ± 5.3 | 43.5 ± 6.3 | 41.9 ± 9.7 |
|  | Frequency (Hz) | 0.33 ± 0.02 | 0.32 ± 0.02 | 0.33 ± 0.03 |
| hM3D | Concentration | 33.5 ± 3.1 | 53.9 ± 6.4<sup>***</sup> | 62.9 ± 5.5<sup>**</sup> |
|  | Frequency | 0.31 ± 0.01 | 0.53 ± 0.02<sup>*</sup> | N.A. |
| hM4D | Concentration | 59.3 ± 5.8 | 32.1 ± 0.4<sup>**</sup> | 19.0 ± 2.3<sup>***#</sup> |
|  | Frequency | 0.30 ± 0.02 | 0.08 ± 0.01<sup>****</sup> | 0.05 ± 0.01<sup>****</sup> |

Data are presented as means ± SEM. $p < 0.05^{**}$ , $p < 0.01$ , $^{***}p < 0.001$ , $^{****}p < 0.0001$ vs vehicle, $^{#}p < 0.05$ vs CNO. N.A., not analyzable.

### Effect of chemogenetic inhibition of VTA-DA on methamphetamine (METH)-enhanced DA regulation in the mOT of anesthetized rats

Based on the findings described above, we investigated how chemogenetic inhibition of VTA-DA neurons impacts METH-induced DA transmission in the mOT of anesthetized rats. In these experiments low and high doses of METH (0.5 and 2.0 mg/kg) were used based on their ability to induce sensitization, place conditioning, and reinstatement consistently^37, 38^ as well as our recent studies on METH effects on DA transmission in the NAc and locomotion^26, 27^. Figure 5 shows representative electrically evoked [DA]_max_ and t_1/2_ changes in the mOT of anesthetized rats in mCherry (left) and hM4D (right) rats 40 minutes after systemic administration of CNO (1.0 mg/kg) followed by METH (0.5 mg/kg or 2.0 mg/kg). METH dose-dependently enhanced both [DA]_max_ and t_1/2_ in mCherry rats, similar to our previous studies in naïve rats^26, 27^. In contrast, the magnitude of METH effects on [DA]_max_ vs. pre-drug control was significantly less in hM4D rats pretreated with CNO compared to mCherry rats (*F*(2,22) = 29.70, *p* < 0.0001). Specifically, at the low dose of METH (0.5 mg/kg), [DA]_max_ did not reach pre-drug control values. At the higher dose (2.0 mg/kg), [DA]_max_ was not significantly different from pre-drug control, however, the t_1/2_ was longer (*F*(2,22) = 29.19, *p* < 0.0001).

**Figure 5:**
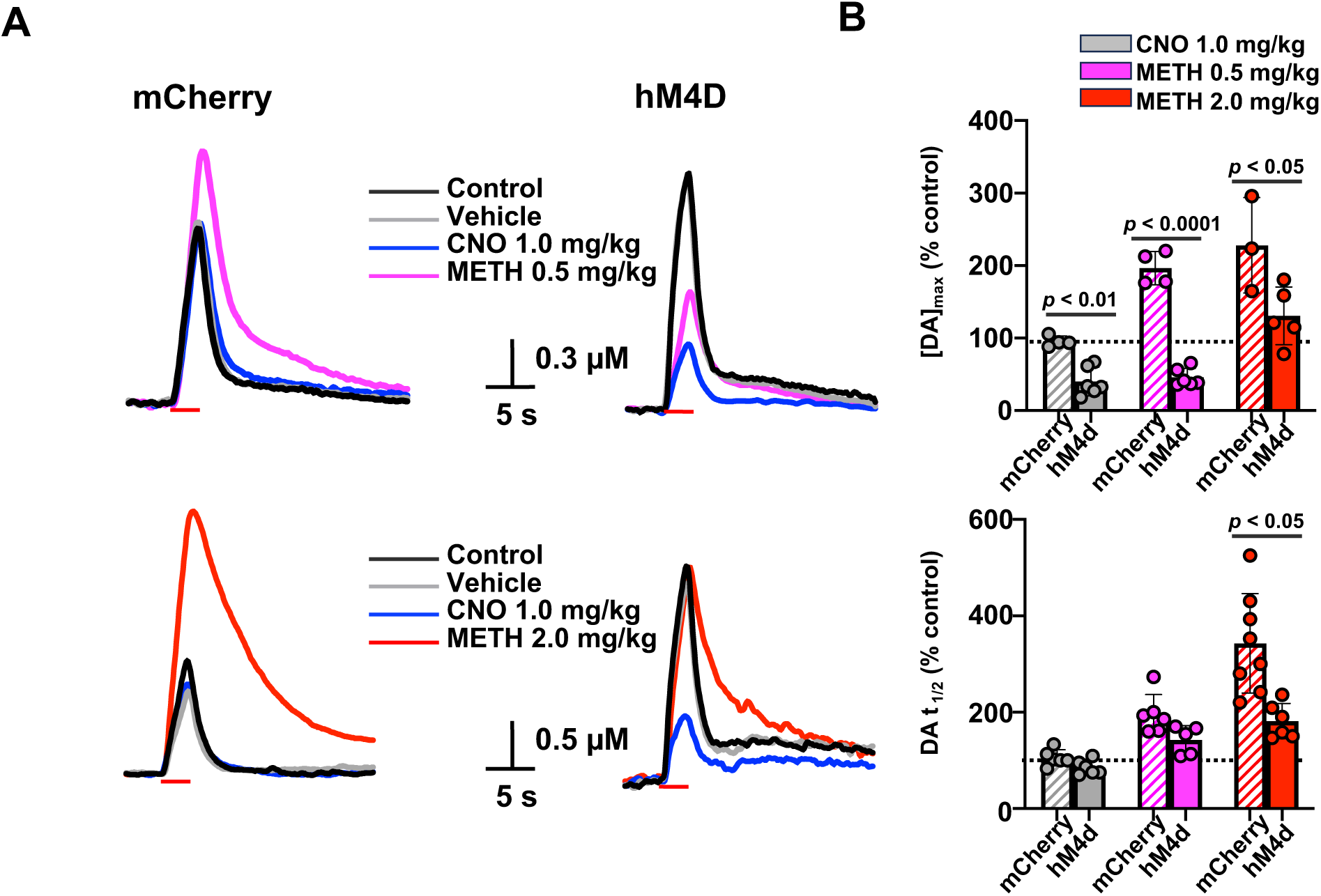
**A.** Representative DA concentration vs time traces recorded in the OT following electrical stimulation of the VTA after vehicle, CNO (1.0 mg/kg) and METH (0.5 mg/kg, top; 2.0 mg/kg, bottom) administered 30 minutes apart in mCherry (left) and hM4D (right) rats. **B.** Average maximal change in electrically evoked [DA]_max_ (top) and t_1/2_ (bottom). Red bars indicate the period of electrical stimulation (20 Hz, 60 pulses). *p < 0.05,**p < 0.01,****p < 0.0001.

### Chemogenetic inhibition of VTA-DA attenuates METH-induced behaviors and DA transmission in the mOT

Naïve (virus-free controls), mCherry, and hM4D rats were administered 1.0 mg/kg CNO 40 min before being subjected to cumulative doses of METH (0.2-2.0 mg/kg) in an open field arena. Figure 6A shows representative locomotor tracks following systemic administration of saline and 0.5 mg/kg METH in mCherry and hM4D rats following CNO administration. Although METH dose-dependently enhanced locomotion (*F*(4, 100) = 26.43, *p* < 0.0001) and stereotypy (*F*(4, 100) = 159.1, *p* < 0.0001) in both groups, hM4D rats demonstrated significantly less locomotion (*F*(2, 100) = 3.841, *p* = 0.0247) and lower stereotypy scores (*F*(4, 100) = 3.413, *p* = 0.0368). However, only at a dose of 0.5 mg/kg METH were these differences significant. mCherry and naïve rats showed nearly identical behavioral responses at all doses tested (Figures 6B and C).

**Figure 6:**
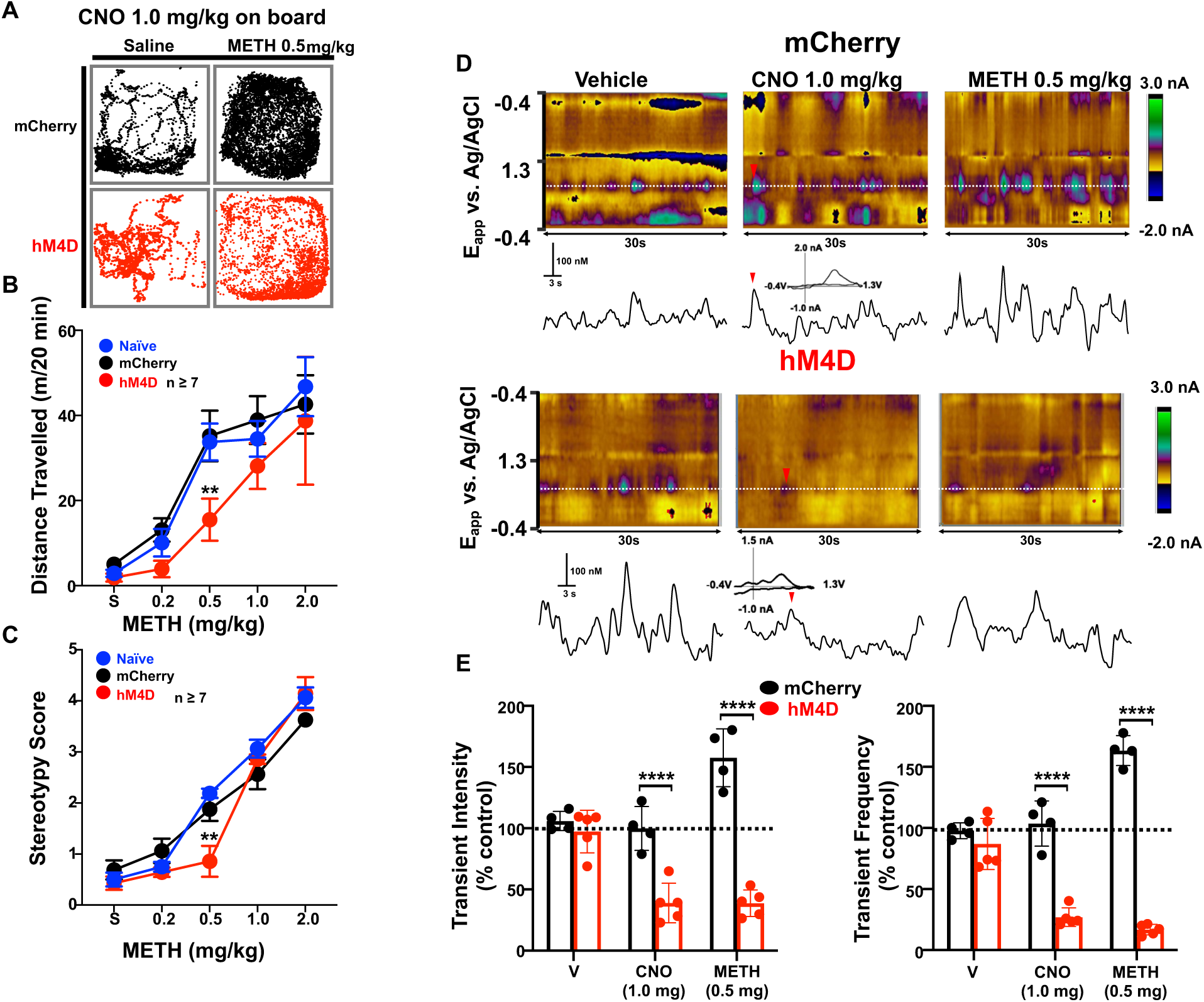
**A.** Representative locomotion tracks in mCherry and inhibitory (hM4D) rats after saline (1.2 mL/kg) and METH (0.5 mg/kg) following CNO (1.0 mg/kg) pretreatment. Changes in locomotion (**B**) and stereotypy (**C**) in naïve (no virus/surgery), mCherry, and hM4D rats after cumulative doses of METH (0.2-2.0 mg/kg) systemically administered 25 minutes apart. **D.** Representative color plots (top) and dopamine (DA) concentration vs time changes (bottom) at the oxidation potential (∼ 0.67 V) for DA (dotted white line) depicting changes in phasic DA following vehicle, CNO (1.0 mg/kg), and METH (0.5 mg/kg) in mCherry and hM4D rats. Inverted red triangles indicate the time when a background subtracted voltammogram was extracted from to identify the analyte. **E.** Average changes from baseline (dotted black line) on transient frequency and intensity in hM4D rats after vehicle, CNO, and METH. ****p < 0.0001.

Next, we tested whether VTA-DA inhibition impacts METH-enhanced naturally occurring DA transmission in the mOT of awake-behaving rats. Figure 6D shows representative color plots and DA concentration vs time traces in mCherry and hM4D rats 40 minutes after vehicle, CNO (1.0 mg/kg), and METH (0.5 mg/kg) sequentially. Voltammograms from a single time point (indicated by red triangles) identify the signal as catecholamine. In these experiments the low dose of METH (0.5 mg/kg) was selected because it is similar to doses used recreationally ^26^. In addition, higher doses of METH can exert non-exocytotic effects on VTA-DA neurons that may supersede DREADD-induced inhibitory effects^26^. Chemogenetic inhibition of VTA-DA significantly reduced METH-induced increases in both the frequency (*F*(2,8) = 49.82, *p* < 0.0001) and intensity (*F*(2,8) = 49.82, *p* < 0.0001) of DA transients in hM4D rats (Figure 6E).

### Activation and inhibition of VTA-DA neurons impact reward-learning

The rewarding/aversive effects of chemogenetic excitation/inhibition of VTA-DA using a conditioned place preference (CPP) paradigm were investigated^39, 40^. Following 10 days of conditioning, chemogenetic activation of VTA-DA neurons with CNO (1.0 mg/kg, IP) induced a strong place preference for the CNO-paired compartment (*F*(2,20) = 3.733, *p* = 0.0419, Figure 7A). However, mCherry and hM4D rats did not show any change in preference following CNO pretreatment. The effects of chemogenetic inhibition of VTA-DA on METH-induced CPP was also investigated. In this study, either mCherry or hM4D rats were given CNO (1.0 mg/kg) or vehicle in their home cages. 40 minutes later, rats were administered either METH (0.5 mg/kg) after CNO or saline (1.2 mL/kg) after vehicle in the CPP chambers. Remarkably, after 10 days of conditioning, hM4D rats pretreated with CNO show avoidance of the METH-paired chamber whereas mCherry rats exhibited a very strong place preference towards the METH-paired chamber (*t*(20) = 2.674, *p* = 0.0073, Figure 7B).

**Figure 7:**
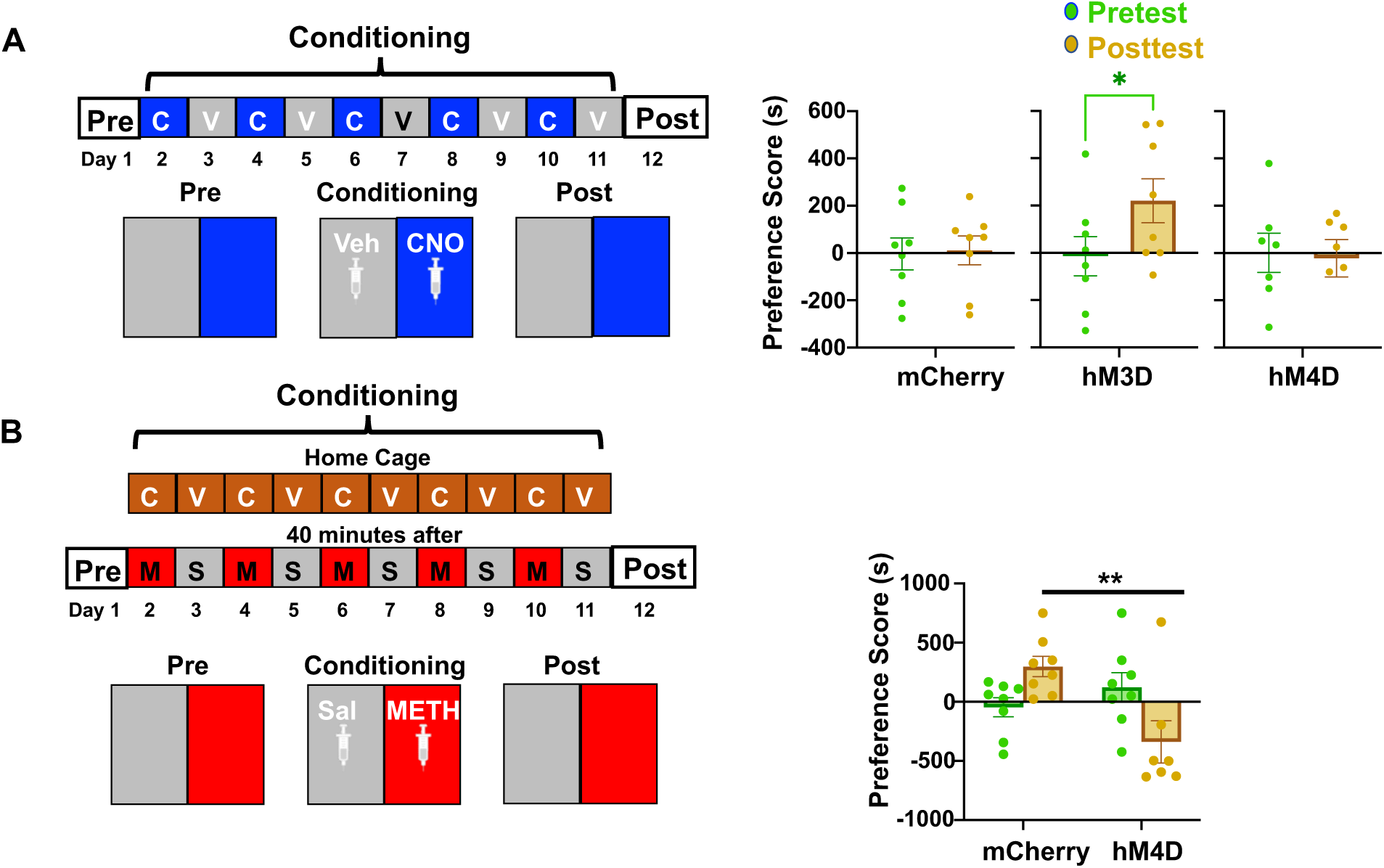
**A.** Change in preference score in seconds (defined by the time spent in CNO-paired compartment during test session minus time spent in CNO-paired compartment during pre-test session) following 10 days of conditioning (CNO 1.0 mg/kg, even days; vehicle, 1.2 mL/kg, odd days) of in mCherry, hM3D, and hM4D rats. Effect of chemogenetic inhibition on METH-induced CPP following Vehicle and CNO-paired METH-pretreatment on alternate days **(B)**. Experiment design schematic shown to the left of each graph. CNO (1.0 mg/kg); S: Saline, V: Vehicle; M: METH (0.5 mg/kg). *p < 0.05, **p < 0.01. n = 7-10 rats/group.

## Discussion

The OT is increasingly recognized for its roles in processing sensory and reward related information, however, understanding the impact of DA transmission in the OT in encoding such modalities has been hindered by anatomical and technical difficulties in selectively controlling and determining local DA signaling. In this study, we investigated VTA-DA transmission in the mOT, a key mediator of reward and drug use disorders^16^. Herein, we demonstrate for the first time how chemogenetic modulation of VTA-DA neurons attenuates or potentiates extracellular terminal DA transmission in the mOT and the associated behavioral outputs in wild-type rats after establishing and optimizing chemogenetic experimental procedures (e.g. doses of CNO, effect times) through integration with *in vivo* FSCV, a neurochemical sensing technique.

### CNO dose-dependently modulates VTA-DA transmission in the OT of anesthetized rats

Many studies have used DREADDs to elucidate the different roles of the mesolimbic DA system in learning, locomotion, and drug seeking, however, little is known about how chemogenetic modulation of DA transmission to produce behavioral effects^28, 41–43^ or how different doses of CNO affect these outcomes. The effects of excitatory (hM3D) and inhibitory (hM4D) DREADDs are based on activation of the G_q_ and G_i_ pathways, respectively^44^. Activation of hM3D increases mobilization of intracellular calcium stores, which leads to the depolarization of neurons and increases in burst firing^45^. In contrast, hM4D activation enhances GIRK channel activity and decreases cAMP production, resulting in hyperpolarization of the membrane potential and a reduction in firing rate^46, 47^. Accordingly, a recent *ex vivo* study showed that bath application of CNO (1 µM) increases the firing rate of VTA-DA neurons expressing hM3D DREADDs as well as electrically evoked DA in the NAc of rat brain slices^48^. Stimulation of hM4D expressing VTA-DA terminals in the NAc also showed a decrease in electrically evoked DA under the same conditions^48^.

Our neurochemical data show that 0.3 mg/kg of CNO is sufficient to activate both excitatory and inhibitory DREADDs that enhance and decrease electrically evoked DA as well as its rate of release in the mOT, respectively. 1.0 mg/kg CNO induces a maximal increase or decrease in extracellular DA transmission in the mOT (Figures 2-4). Importantly, our results highlight that CNO effects on DA transmission are maximized within 30-40 minutes after systemic administration and last for at least 1 hour for both anesthetized and awake behaving rats (Supplementary Figure 2 and Figure 4), which is similar to previous neurochemical (e.g. DA and oxytocin) and behavioral studies^25, 49^. It has also been reported that CNO (0.3 - 1.0 mg/kg) can activate/inhibit DA neurons for 6-10 hours and alters locomotion up to 11 hours after CNO administration in rodents^3^.

The effects of chemogenetic excitation and inhibition of VTA-DA neurons on electrically evoked DA release in the mOT of anesthetized rats resemble those observed following administration of DA D2 autoreceptor antagonists and agonists, respectively. Administration of raclopride (D2 antagonist) significantly potentiates DA release and increases its slope in anesthetized mCherry rats, similar to previous studies in naïve rats (Figure 3)^9, 30^. In contrast, the D2 receptor agonist (quinpirole) causes a significant decrease in electrically evoked DA in brain slices (∼80 – 90 %) and anesthetized rats (∼60 %), depending on the electrical stimulation frequency and duration^50–52^. In this study, DA release evoked by electrical stimulation (20 Hz, 60 pulses) was decreased ∼ 70% as well as the slope of release following CNO administration in hM4D rats. The latency for DA release was also delayed and not observable at the beginning of the stimulation (shorter stimulation trains, ≤ 10 pulses)(Figure 3). Therefore, to accurately characterize DA regulation, higher stimulation pulse numbers (60 pulses) were used to generate reproducible release. In agreement with this result, an electrophysiology study in GABA neurons expressing hM4D DREADDs showed that CNO completely silences neuronal activity during low current injection but only partially attenuates it during high current injection^53^. These results additionally support the hypothesis that in the absence of electrical stimulation or at low stimulation frequencies/durations, DA release may not be detectable following chemogenetic inhibition of VTA-DA neurons (signal to noise ratio, S/N<5). The attenuated DA release was not reversed by raclopride, but the DAT inhibitor, GBR 12909, relatively increased evoked DA concentration like mCherry rats (Figure 3). Our previous studies have shown that in naïve rats, naturally occurring phasic DA release is observed in the NAc and OT after D2 antagonism and inhibition of the DAT^9, 13, 30, 35^. Consistent with previous studies, mCherry rats exhibited robust phasic DA transients in the mOT following combined administration of the D2 antagonist and DAT inhibitor, while CNO (1.0 mg/kg) administration had no effect on DA transients. Conversely, no naturally occurring transients were observed in hM4D rats that were administered CNO (1.0 mg/kg) under identical experimental conditions (Figure 3D). Therefore, these results agree with previous electrophysiology studies and confirm that chemogenetic activation and inhibition of VTA-DA cell firing modulates DA release but with little impact on DA uptake.

Importantly, there were no significant effects of CNO on both electrically evoked and naturally occurring phasic DA in anesthetized and awake behaving mCherry rats at any CNO dose (< 3.0 mg/kg) tested (Figures 2 & 4 and Supplementary Figure 3). It is noteworthy that such doses (lower than 3.0 mg/kg) administered systemically appear to be pharmacologically inert for rats in most behavioral models, particularly those with a heavy mesolimbic influence^23, 54^. Increasing evidence has suggested that CNO can be reverse-metabolized into the psychoactive compounds clozapine or N-Desmethylclozapine, which can produce off-target effects on central nervous system function and basal behaviors by binding to endogenous receptors^55–57^. Moreover, clozapine can bind to D2 receptors and directly impact DA transmission^28, 29, 36, 58^. However, many studies continue to use high doses of CNO (≥ 5 mg/kg, IP) that can impair DA D2/3 and serotonin 5HT_2A_ receptor occupancy and impact the regulation of many neurochemicals and behaviors^59^. For example, high CNO doses (5.0-8.0 mg/kg) reduce glutamate levels in the striatum^60^ as well as disrupt behavioral responses (e.g. acoustic startle reflex)^61^. Although these effects may be related to elevated levels of clozapine in the brain, some evidence has been shown that the back-conversion to clozapine is negligible at CNO doses less than 5.0 mg/kg in rats, however these findings remain controversial^55, 57^. It is also worth noting that the metabolism of CNO to other compounds may be species-dependent, as rats have been shown to exhibit lower levels of clozapine in the brain following systemic administration of equivalent doses of CNO to mice^62^. Taken together, our results and previous studies suggest that 1.0 mg/kg CNO is sufficient to modulate VTA-DA in DREADD expressing rats with minimal effects on DA regulation in control rats.

### The functional link between distinct behavioral outputs and mesolimbic DA transmission in the OT

It is generally known that VTA-DA plays an important role in fundamental behaviors necessary for survival including the initiation of movement and locomotion^2^. Systemic administration of D1 antagonists reduce spontaneous locomotion in rats^63^ whereas microinfusions of DA or D1 receptor agonists into the ventral striatum enhance locomotor activity^64, 65^. In contrast, similar experiments utilizing local injection of D2 agonists have been shown to reduce spontaneous locomotion^66^. Our behavioral data show that CNO dose-dependently enhanced locomotion and rearing in hM3D rats, but 3.0 mg/kg CNO did not significantly increase the spontaneous hyperlocomotion more than 1.0 mg/kg (Figure 4). Activation of VTA-DA with CNO causes an increase in the concentration of naturally occurring DA transients which coincides with enhanced horizontal and vertical locomotion in hM3D rats (Figure 4, Table 1). Importantly, at a higher dose of CNO (1.0 mg/kg), phasic transients cannot be accurately quantified due to the presence of overlapping DA signals and technical constraints. More specifically, besides the anatomical challenges (e.g., misplacement of neurochemical sensors in the OT and off-target virus infusions in the VTA) of these experiments, hM3D rats showed such intense hyperactivity that it often resulted in damage to the brittle carbon-fiber microelectrode and electrical noise during neurochemical recording. Other behaviors, including jumping and darting, often led to attrition due to the cranial implant site becoming compromised. Therefore, the success ratio of hM3D awake-behaving animal experiments coupled with FSCV recording was incredibly low ∼ 20 % compared to hM4D experiments (∼ 60% success ratio). Additionally, while we cannot ignore the possibility of some virus spillover into the substantia nigra (SN), which may impact chemogenetically enhanced DA release and behavior, we minimized viral infusion volumes to limit DREADD expression to the VTA as much as possible (Figure 1). In addition, the locomotor changes observed are in agreement with previous studies utilizing both optogenetics^67^ and chemogenetics^28, 68^ showing that VTA-DA but not SN-DA neurons are involved in inducing locomotor hyperactivity^28, 29^. Instead, activation of SN-DA neurons played a larger role in other aspects of locomotion including motor coordination as well as orientation and acceleration^28^. The locomotor data from hM4D rats likely reflect a floor effect as we did not observe a significant reduction in behavior. Despite a slight but not significant decrease in locomotion in mCherry rats, this was likely due to a decrease in exploratory behavior over time during the 160-minute session.

The results from place conditioning experiments additionally demonstrate that chemogenetic activation of VTA-DA is rewarding (Figure 7A), consistent with previous reports in transgenic mice^39, 40^. Recordings of VTA-DA transmission in the mOT of the wild-type rats and behaviors suggest that both an enhanced concentration and frequency of naturally occurring phasic DA transmission in the mOT may be implicated in distinct locomotor and reward outputs. In contrast, hM4D rats showed no significant locomotor changes and alteration of place preference (Figure 6 & 7A), despite phasic DA release being significantly attenuated in the mOT of awake-behaving rats, similar to electrically evoked data from anesthetized rats. While a previous study showed that chemogenetic inhibition of VTA-DA decreases locomotion ^40^, our open field testing was performed during the rats’ light cycle while the former study occurred during the dark cycle, during when mice and rats are generally more active. Therefore, the inhibitory effect of VTA-DA on locomotion may be stronger during their dark cycle^69^. Additionally, another study showed a significant reduction of locomotion in hM4D mice following CNO administration, however, this may be more pronounced in mice due to a greater baseline locomotion level^68^.

### Chemogenetic inhibition of VTA-DA neurons attenuates METH-enhanced DA transmission in the OT and reward seeking behaviors

Chemogenetics has many distinct advantages over conventional techniques for transiently modulating neuronal activity including its long-lasting effects (minutes to days) and ability to selectively target neuron subtypes. This is particularly beneficial for studying drug use disorders, especially for psychostimulants that have a long half-life in the body and enables parsing out specific neurochemical components of distinct brain regions that drive different drug-induced behaviors. In both anesthetized and awake-behaving animal experiments, we demonstrate for the first time, that chemogenetic inhibition of VTA-DA neurons suppresses both METH-enhanced electrically evoked and naturally occurring DA transmission in the OT as well as METH-induced increases in locomotion (Figure 5 & 6). METH (0.5 mg/kg) did not increase electrically evoked DA above pre-CNO control nor restore the CNO-suppressed phasic DA transients in the OT after chemogenetic inhibition of VTA-DA neurons. Furthermore, following CNO, hM4D rats displayed a reduction in METH-induced hyperlocomotion and stereotypy behaviors at lower METH doses (< 1.0 mg/kg) compared to virus-free naïve and mCherry rats pretreated with CNO (Figure 6). This is in agreement with similar studies showing chemogenetic inhibition of VTA-DA with CNO (2.0 mg/kg) in transgenic mice reduces cocaine-induced hyperlocomotion for at least 1 hour^40^. These results also suggest that VTA-DA and its transmission are necessary for inducing specific changes in the locomotor effects of psychostimulants. In contrast, a higher dose of METH (2.0 mg/kg) causes not only [DA]_max_ to return to pre-drug control values yet the t_1/2_ was still enhanced greater than the control values (Figure 5). This may be due to non-exocytotic effects (e.g. reverse transport) or other mechanisms enhancing extracellular DA concentration as there were no significant METH-enhanced behavioral differences between hM4D and mCherry rats at this dose^26^. Additionally, as evidenced by our pharmacological study (Figure 3), chemogenetic inhibition of VTA-DA had very little impact on DA reuptake. Therefore, METH, a monoamine transporter inhibitor, can still dose-dependently enhance extracellular DA concentrations. It is also noteworthy that the behaviors following chemogenetic excitation of VTA-DA are very similar to those reported by microinfusions of psychostimulants such as amphetamine and cocaine into the OT, which potentiates both horizontal and vertical locomotion^65^, but do not produce the focused stereotypies (in place biting, sniffing, chewing) often induced by high doses of amphetamines or cocaine^70^. This further suggests that the locomotor changes seen were due to enhanced VTA-DA in the ventral striatum.

In addition to locomotion, chemogenetic inhibition of VTA-DA also prevented the acquisition of METH-induced reward. Surprisingly, CNO-pretreatment before METH in hM4D rats led to avoidance for the METH-paired compartment. These results suggest that despite a lack of change in place preference following chemogenetic inhibition of VTA-DA alone (Figure 7A), inhibition of VTA-DA neurons was strong enough to attenuate the rewarding effects of low doses of METH, which may be due to an imbalance between METH-induced alterations of the reward (dopamine) and stress (norepinephrine) systems. Importantly, METH (compared to other psychostimulants such as cocaine) has approximately 2.5 times higher affinity for the norepinephrine transporter than DAT, enhancing extracellular norepinephrine concentrations via decreased uptake more than DA at the same dose^27, 71^. Thus there may be METH-enhanced extracellular norepinephrine concentrations that increase aversion while the METH-induced enhancement of DA transmission in limbic areas, including the OT, is attenuated by inhibition of VTA-DA neurons. Interestingly, a similar study showed that chemogenetic inhibition of D1 receptor expressing GABAergic medium spiny neurons in the NAc is sufficient to prevent the expression of cocaine (20 mg/kg) induced-CPP^53^. Therefore, inhibition of ventral striatal region-specific VTA-DA innervation of the OT and/or NAc may distinctly mediate the rewarding properties of psychostimulants. Our findings along with previous studies highlight that chemogenetic inhibition of VTA-DA is able to attenuate METH effects on DA transmission in the OT, which may in turn lower the rewarding effects of METH and induce aversion. While VTA-DA signaling in the NAc has been extensively studied, our and previous research suggests DA in the OT also significantly contributes to both the rewarding effects of psychostimulants and modulating psychostimulant-induced reward^16, 72^. However, future studies integrating specific targeting of the VTA-DA projection neurons to the OT (VTA-DA→OT pathway) will be conducted to further test whether a functional link between VTA-DA transmission in the OT is necessary for modulating the reinforcing effects of reward and drug use disorders.

### Conclusion

Through use of neurochemical sensing and anesthetized/awake behaving animal experiments we optimized and demonstrated for the first time how the ligand CNO dose-dependently modulates dopamine transmission in the mOT and behaviors via chemogenetic manipulations (activation/inhibition) of VTA-DA neurons. These findings highlight a correlation between enhanced DA transmission in the mOT and positive behaviors (e.g. enhanced locomotion/exploration and place preference). In contrast, inhibition of VTA-DA signaling may decrease the salience of otherwise rewarding stimuli via suppression of DA transmission in the mOT and attenuate expression of such behaviors. Taken together, these results provide the framework for future behavioral and neurochemical studies by elucidating how DREADDs modulate local DA transmission and associated behavioral changes as well as further highlight the role of the OT DA in addiction research.

## Methods

### Experimental Subjects

Wild-type, male Sprague-Dawley rats (n = 144) (280 to 350 g, aged ∼8 weeks at the time of arrival) were obtained from Charles River Laboratories (Wilmington, MA, USA). Sample sizes were determined based on earlier studies and on our preliminary data to main a power of 0.8.

Rats were initially pair-housed in a polycarbonate cage with a wire lid holding rat chow within a humidity and temperature-controlled environment with a 12:12h light-dark cycle (lights on at 07:00h). Following stereotaxic surgery, rats were housed individually in clean cages to prevent damage to the surgical site. Food and water were available *ad libitum*. Water was delivered through an animal watering system via a small opening in each cage. All procedures related to handling and caring for animals were performed in accordance with the ‘Guide for Care and Use of Laboratory Animals’ (8th Edition, 2011, US National Research Council) and were approved by the Institutional Animal Care and Use Committee of the University at Buffalo (Protocol CLS08085Y).

### Stereotaxic surgeries for virus infusion and DA recording

#### (1) Viral mediated gene transfer

In order to exclusively excite or inhibit VTA-DA transmission in the OT, excitatory (hM3D) and inhibitory (hM4D) DREADDs, respectively, were selectively expressed in VTA-DA neurons. Transgene expression was achieved in wild-type rats through use of a combinatorial viral targeting system. To obtain specificity for DA neurons, the targeting virus (TH2.6-iCre-AAV2/10) drove improved Cre-recombinase (iCre) expression in tyrosine hydroxylase (TH) expressing neurons^23^. The effector viruses used (EF1α-DIO-hM3dq-mCherry-AAV2/10 and EF1α-DIO-hM4Di-mCherry-AAV2/10) carried the DREADD transgene in a Cre dependent double-floxed inverted open reading frame (DIO) under control of the human elongation factor-1 alpha (EF1α) promoter^23^. Therefore, only TH+ neurons expressing iCre expressed the DREADDs. mCherry controls were infused with a virus encoding for a red fluorescent protein (mCherry, EF1α-DIO-mCherry). All viruses were mixed with 1 part targeting virus to 2 parts effector virus and tittered to approximately 10^13^ vector genome copies per mL and were generated in-house.

Rats were injected with ketamine (85 mg/kg) and xylazine (10 mg/kg) and stereotaxic surgery was performed based on previous studies^13, 35^. Rats were placed in a stereotaxic frame (David Kopf Instruments, Tujunga, CA, USA) and a central incision was made to expose the skull surface. Anteroposterior (AP), mediolateral (ML) and dorsoventral (DV) positions were referenced from Bregma^73^. A microsyringe (26g, 10 μL volume, Hamilton Company, Reno, NV, USA, Model# 701N) was slowly lowered into the VTA (AP: -5.2 mm, ML: -0.8 mm, DV: -7.8 mm) at a rate of 1 mm/min. Then, 500 nL of virus was injected (100 nL/min) through the Hamilton syringe connected to a syringe pump with stereotaxic holder (KD Scientific). To ensure complete diffusion of virus, the syringe was left in place for 10 minutes after the injection and then slowly retracted. Injections were performed bilaterally. 4 weeks were allowed for transgene expression and animal recovery.

#### (2) Voltammetry surgery

##### Survival voltammetry surgery

4 weeks after virus infusion, rats were injected with ketamine and xylazine and stereotaxic surgeries for electrochemical measurements were performed as described previously^26, 74, 75^. Burr holes (1 mm diameter) were drilled in the skull for implanting screws and electrodes. A guide cannula (Bioanalytical Systems, West Lafayette, Illinois) for loading a micromanipulator containing a fresh carbon-fiber microelectrode (neurochemical sensor) was implanted above the OT (AP: +1.8 mm, ML: 0.9 mm). An untwisted bipolar stainless-steel stimulating electrode (Plastics One, Roanoke, VA, USA) insulated to the tips (0.2 mm diameter, separated by ∼1.0 mm) was implanted into the VTA (AP –5.2 mm, ML 1.0 mm, DV ∼ -8.0-9.0 mm). A Ag/AgCl reference electrode was implanted on the contralateral cortex. All items were secured with stainless steel skull screws and dental cement (Lang Dental, #1234). Animals were allowed 1 week to recover before neurochemical and behavioral recordings.

##### Non-survival anesthetized voltammetry surgery

4 weeks after virus infusion, rats were anesthetized with an intraperitoneal (IP) injection of urethane (1.5 g/kg) and placed in a stereotaxic frame. Holes were drilled for the insertion of neurochemical sensors, reference and stimulating electrodes^13, 26^. Urethane was selected as the appropriate anesthetic because tonic neuronal firing is not reduced nor is electrically evoked DA, such as with ketamine anesthesia^76, 77^. Upon puncturing and removal of the dura, a neurochemical sensor was vertically implanted into the *m*OT (AP: +1.8 mm, ML: +0.9 mm, DV: ∼ -8.0 to 8.5 mm) and a stimulating electrode was implanted into the VTA (AP: –5.2 mm, ML: 1.0 mm, DV: ∼ -8.0-9.0 mm) on the right hemisphere. A Ag/AgCl reference electrode was implanted into the contralateral hemisphere and secured with a skull screw.

#### (3) Voltammetric DA recording procedures

Glass-encased carbon fiber and Ag/AgCl reference electrodes were prepared as previously described^13, 26, 75, 78^. T-650 untreated carbon fibers (Cytec Industries Inc., Greenville, SC, USA) with 7 µm in nominal diameter were used and cut to an exposed length of 75-100 μm^34, 35^. For FSCV recordings, TH-1 software (University of North Carolina Department of Chemistry Electronic Shop)^30, 79^ was used to apply a triangular waveform (-0.4 V to +1.3 V and back to -0.4 V, 400 V/s, 100 ms intervals) to the carbon-fiber microelectrode. The triangular waveform was low-pass filtered at 2 kHz. Background-subtracted cyclic voltammograms were obtained by performing a digital subtraction of voltammograms collected during baseline recording before electrical stimulation trains and administration of pharmacological agents. These voltammograms were used to identify the catecholamines (DA and norepinephrine), as they are distinct from their metabolites and other neurochemicals^30, 80, 81^. Voltammetric responses were viewed as color plots with the abscissa as potential, the ordinate as acquisition time, and the current encoded in false color. Current was converted to concentration based on calibration curves with known concentrations of DA (10.5 ± 0.4 pA/(μM·μm^2^))^9, 13, 26^. At the end of all experiments, electrode locations in the mOT were confirmed by electrolytic lesion^9, 30^.

#### (4) Electrical stimulation

For anesthetized animal experiments, electrical stimulation of the VTA consisted of biphasic square wave pulses (300 µA, 2 ms/phase) that were computer-generated with a 6711 PCI card (National Instruments, Austin, TX, USA) and were optically isolated from the electrochemical system (NL 800A, Neurolog, Digital Ltd., Hertfordshire, UK). The stimulus parameters were held constant at 20 Hz, 60 pulses during experiments^9, 30, 35^. Stimulation trains were repeated every 10 minutes, which allows for reproducible DA release^9, 35, 82^. For awake-behaving experiments, stimulus intensity (80-150 µA) and pulse numbers were varied to evoke different concentrations of DA and pH in the OT at the end of experiments in order to construct a training set for chemometric analysis^75, 83^.

### Immunohistochemistry and histology

Following behavioral or FSCV recordings, all rats were deeply anesthetized with sodium pentobarbital (487.5 mg/kg) and transcardially perfused with 500 mL of phosphate buffered saline (PBS, 4°C) followed by 350 mL of 10% buffered formalin phosphate (4°C). For FSCV experiments, neurochemical sensor placement was verified by applying constant current (20 μA for 10 s) directly to the electrode^9^ before the perfusion. Brains were extracted from the skull, postfixed for 12 hours in 10% buffered formalin phosphate at 4°C and then cryoprotected in 30% sucrose in 0.1 M PBS for 72 hours. Brains were sectioned on a freezing microtome at 35 and stored in PBS containing 0.02% sodium azide until free-floating sections were processed for immunofluorescence (IF). Sections were washed in 0.1 M PBS, pH 7.4 (3 x 5 minutes) to remove residual sodium azide, then permeabilized and blocked for 2 hours in 5 % normal goat serum (Vector, S-1000)/0.5 % Triton-s/0.1 M PBS. They were then directly placed in the primary antiserum for 2 hours at room temperature and then incubated overnight at 4°C. Primary antibodies consisted of a mouse anti-tyrosine hydroxylase (ImmunoStar #22941, 1:1000) and a rabbit anti-RFP (Rockland #600-406-379, 1:10,000) diluted in PBS + 0.5% triton X-100. The next day, sections were rinsed in PBS (3 x 10 minutes) and then incubated with secondary antibodies consisting of Alexa 488 goat anti-mouse (Invitrogen #A11029, 1:1000) and Alexa 555 goat anti-rabbit (Invitrogen #A21429, 1:1000) for 2 hours at room temperature. Sections were rinsed again in PBS (3 x 10 minutes), mounted on slides and coverslipped using Prolong Diamond mounting media (Life Technologies, USA). Slides were visualized on an Andor Dragonfly spinning disk confocal microscope at the Optical Imaging and Analysis Facility, School of Dental Medicine, State University of New York at Buffalo. For verification of electrode placement, sections containing the electrolytic lesion were mounted on slides and viewed under a stereo microscope with blue LED illumination (AMScope) microscope.

### Behaving FSCV

Rats were habituated for two days (1 hour/day) to a custom-built open field arena (12” x 12” x 24”) with a video camera mounted above and housed within a Faraday Cage (to reduce electrical noise). On the recording day, rats were implanted with a detachable micromanipulator directly above the NAc, and the neurochemical sensor was lowered 6.0 mm below the skull surface. A bipolar stimulating electrode was implanted in the VTA to deliver electrical stimulation (60 Hz, 24 pulses) to the DA cell bodies. Generally, a stimulating electrode is not required to place a neurochemical sensor in the NAc since the naturally occurring phasic DA transients are clearly observed between ∼6.0 - 7.5 mm below the skull. However, due to the anatomical challenge of properly placing the sensor in the relatively thin OT, electrical stimulation is necessary to map the distribution pattern of DA release and the boundary between the NAc and OT^9, 13^. Once a stable signal was found, rats were given vehicle (1.2 mL/kg, IP) followed by cumulative doses of CNO (0.3 & 1.0 mg/kg, IP). For METH experiments, rats were administered vehicle and CNO (1.0 mg/kg, IP) before METH (0.5 mg/kg, IP), All drugs were administered 40 minutes apart. Following FSCV recordings, rat behavior was blindly analyzed by scorers naïve to the experiment conditions. Crossovers (the number of times a rat crosses a quadrant in the apparatus) and rearing were manually counted during the first minute of every 5-minute bin^26, 84^.

### Behavioral Measures

#### (1) Open field testing

4 weeks after virus infusions, rats were habituated for 2 consecutive days (1 hour/day) to a 16” x 16” x 16” open field arena (E63-20, Harvard Apparatus, Holliston, MA). On day 3, rats were subjected to a 160-minute recording session (4 x 40 minute components) where a CNO locomotor dose-response curve was generated using a cumulative dosing procedure^38^. Rats were injected with vehicle (1.2 mL/kg, IP) followed by cumulative doses of CNO (0.3-3.0 mg/kg, IP). Likewise, METH open field testing was performed in a similar fashion. Following 2 days habitation, rats were administered CNO (1.0 mg/kg, IP) in their home cages. 40 minutes later, rats were subjected to a 125-minute recording session (five 25-minute components) where a METH dose-response curve was generated. Rats were injected with saline (1.2 mL/kg, IP) followed by escalating doses of METH (0.2-2.0 mg/kg, IP)^26^. Locomotion, stereotypy, and rearing counts were measured in 5-minute bins; however, the first 5-minutes of each dose exposure was excluded from analysis due to hyperactivity from injection and touching. Behavior was recorded with a USB webcam (SMC02, Stopmotion Explosion) interfaced with a PC and analyzed using commercially available video tracking software (AnyMaze, Stoelting, Wood Dale, IL). Time-stamped coordinates of animal positions were extracted from the video and plotted to visualize changes in horizontal locomotion. Rearing (vertical locomotion) and stereotypy were manually scored from the video recordings (as described above).

#### (2) Conditioned place preference (CPP)

CPP experiments were based on previous studies^39^ and performed in modified ‘Med Associates Two Chamber Place Preference Inserts for Rats’ (ENV-517, Med Associates, St. Albans, VT) with distinct tactile (wire/grid floor) and visual cues (black and white bars/checkers) in each chamber. A split wall of black Plexiglas separated the chamber into two compartments (8.75” L x 8.75” W x 12” H). During conditioning, compartments were fully separated by the split wall (guillotine door inserted). However, on test days, the split wall contained an opening to allow free passage between the two compartments (guillotine door removed). Time in each compartment was recorded by a digital video camcorder (Canon #HF R70) and place preference was scored using AnyMaze. Schematic timelines for each CPP experiment are shown in Figure 7.

##### (2.1) Pretest (Day 1)

Rats were handled (20 minutes/day) and habituated to the testing room (40 minutes/day) for 2 days. On day 1, rats were randomly placed into one of the CPP chambers with the guillotine door removed and allowed to explore the entire arena for 30 minutes in order to verify the absence of preference to either compartment. The first 15 minutes of the pretest session were analyzed to determine initial preference scores. Rats spending more than 75% (time spent > 675 seconds) or less than 25% (time spent < 225 seconds) of the total time in a compartment were not included in CPP experiments. Following pretesting, rats were randomly assigned to be conditioned in one of the two chambers. Rats were counter balanced and a non-biased procedure was used.

##### (2.2) Place-conditioning (Days 2–11)

Experiment 1 (CNO conditioning): During conditioning sessions, rats were administered VEH (1.2 mL/kg, odd days) or CNO (1.0 mg/kg, even days) and confined to the paired chamber for 30 minutes.

Experiment 2 (CNO/METH conditioning): 40 minutes before conditioning, rats were administered VEH (1.2 mL/kg, odd days) or CNO (1.0 mg/kg, even days) in their home cages. Rats were then administered either saline after VEH (1.2 mL/kg, odd days) or METH after CNO (0.5 mg/kg, even days) and placed into the paired chamber for 30 minutes. Rat groups were denoted as CNO-METH and VEH-Saline.

##### (2.3) Post test (Day 12)

On the day after the last conditioning session, post-conditioning preferences were determined. Rats were again allowed to freely move between the two compartments in a 30-minute test session where time spent in each compartment was recorded.

### Drugs and reagents

CNO and (+)-Methamphetamine-HCl were supplied by the National Institutes of Health Drug Supply Program. CNO was dissolved in 1.5 % ethanol in saline solution. All other drugs and chemicals were reagent-quality and purchased from the Sigma-Aldrich (St. Louis, MO, USA). *In vitro* post-calibration of carbon-fiber microelectrodes was performed in a Tris buffer solution (pH 7.4) containing 15 mM Tris, 140 mM NaCl, 3.25 mM KCl, 1.2 mM CaCl_2_, 1.25 mM NaH_2_PO_4_, 1.2 mM MgCl_2_, and 2.0 mM Na_2_SO_4_ in double distilled water^85^. Raclopride-HCl (Sigma-Aldrich, Cat # R121) and (+)-Methamphetamine-HCl were dissolved in sterile saline. GBR 12909 (Sigma-Aldrich, Cat # D052) was dissolved in water and diluted in sterile saline^9^. Immunofluorescence was performed in a phosphate buffer saline (pH 7.4) containing 27 mM NaH_2_PO_4_ (monobasic monohydrate), 77 mM NaH_2_PO_4_ (dibasic anhydrous), and 154 mM NaCl..

### Data analysis

All datasets generated and/or analyzed during the current study are available from the corresponding author on reasonable request. *Neurochemical data*: Clampfit 8.1 as part of pCLAMP 8.1 software package (Axon Instruments, Foster City, CA, USA) was used to analyze all neurochemical data as described earlier^86^. A training set was created in order to extract the DA component from a mixed analyte signal using a principial component regression algorithm^83^. DA transients were considered signals when they were 5 five times the root-mean-square noise level (S/N > 5). DA transients were analyzed for frequency and amplitude every five minutes after administration of drug and averaged across bins using MATLAB with custom scripts written in-house. [DA]_max_ is the maximally evoked DA concentration and t_1/2_ is the time for [DA]_max_ to decay to half of the maximum^86^. Data are represented as mean ± SEM (standard error of the mean) and ‘n’ values indicating the number of rats. Statistical analysis was performed using GraphPad Software (La Jolla, CA). One-way or Two-way ANOVA with Fisher’s LSD for multiple comparisons or a two-tailed student’s t-test were used to analyze drug-effect data. A mixed model ANOVA with Dunnett’s multiple comparisons test was used to analyze time course CNO effect data. *p* < 0.05 was regarded as statistically significant for all experiments. Rats that showed damage to the sensor or excessive noise were excluded from neurochemical analysis at a specific dose(s) but were still included in behavior analysis. However, any data obtained from the misplacements of the electrodes was discarded.

#### Immunofluorescence

Co-expression of TH and DREADD-mCherry was determined by collecting Z-stacks (approximately 25-35, 685 nm optical slices) from several fields in the VTA and examining them using ImageJ software. mCherry+ cell bodies were carefully examined in each z-stack to determine if a TH signal was also present. *Statistics:* Mann-Whitney U test was used to analyze co-expression of TH and DREADD-mCherry. Rats that did not show DREADD-mCherry expression were excluded from analysis.

#### Behavioral analysis: Locomotion

Locomotor dose-response curves were analyzed by two-way repeated measures ANOVA with Fisher’s LSD for multiple comparisons. Conditioned Place Preference (*CPP)*: The magnitude of place preference was presented as preference scores, which were defined as the time spent in drug-associated compartment during test day minus time spent in the drug-associated compartment during pretest day. Data were analyzed using one-way ANOVA followed by Bonferroni’s post hoc test to determine the effects of drug-induced CPP.

## Abbreviations

AP: anteroposterior
DA: dopamine
DAT: dopamine transporter
DREADDs: Designer Receptors Exclusively Activated by Designer Drugs
DV: dorsoventral
FSCV: fast-scan cyclic voltammetry
IP: intraperitoneal
ML: mediolateral
OT: olfactory tubercle
TH: tyrosine hydroxylase
VTA: ventral tegmental area

## Acknowledgements

This work was funded by NIH/NIDA (DA056547 & DA0452840 and the SUNY Seed Grant (RFP#23-01-RSG). We gratefully acknowledge the support from the NIDA drug supply program. Special thanks to Andrew McCall at the University at Buffalo Optical Imaging and Analysis Facility as well as Michael Fletcher, John Franke, and Brian Koyn at the Health Science Instrument Shop for assistance with image analysis and instrument fabrication, respectively.

## Author Contributions

R.V.B and J.P. conceived the present study and designed experiments. R.V.B and R.C.P performed experiments and analyzed data. R.V.B and J.P. wrote the manuscript. C.E.B. contributed viruses and advised with behavioral experiments. J.P. acquired funding and supervised the study.

## Notes

### Competing Interest Statement

The authors have declared no competing interest.

### Summary of Updates

This revision fixes an error in a behavior figure and removes some confusing wording to better explain experiment design

