## Supplementary Figures for "Decoding mesolimbic dopamine transmission in the olfactory tubercle and its contribution to methamphetamine responses through neurochemical sensing and chemogenetics"

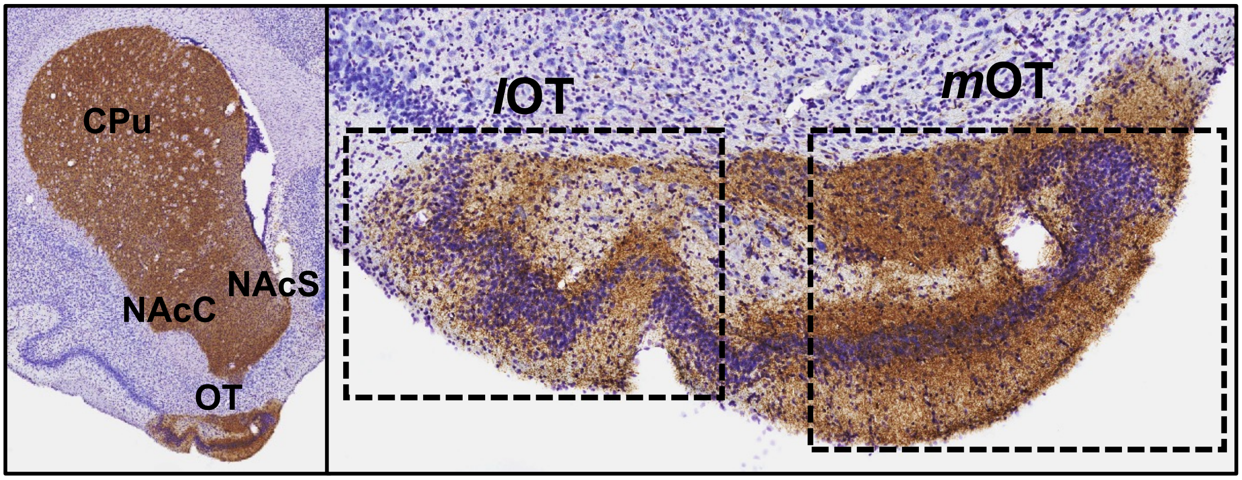


**Supplementary Figure 1**: Tyrosine hydroxylase (TH) expression throughout the ventral striatum. **Left:** coronal section of the ventral striatum. **Right:** magnification of the OT subregion of the ventral striatum. Brown color indicates TH expression, marker for catecholamines (DA). CPu: Caudate Putamen; NAcC: Nucleus Accumbens Core; NAcS: Nucleus Accumbens Shell; mOT: Medial Olfactory Tubercle; lOT: Lateral Olfactory Tubercle

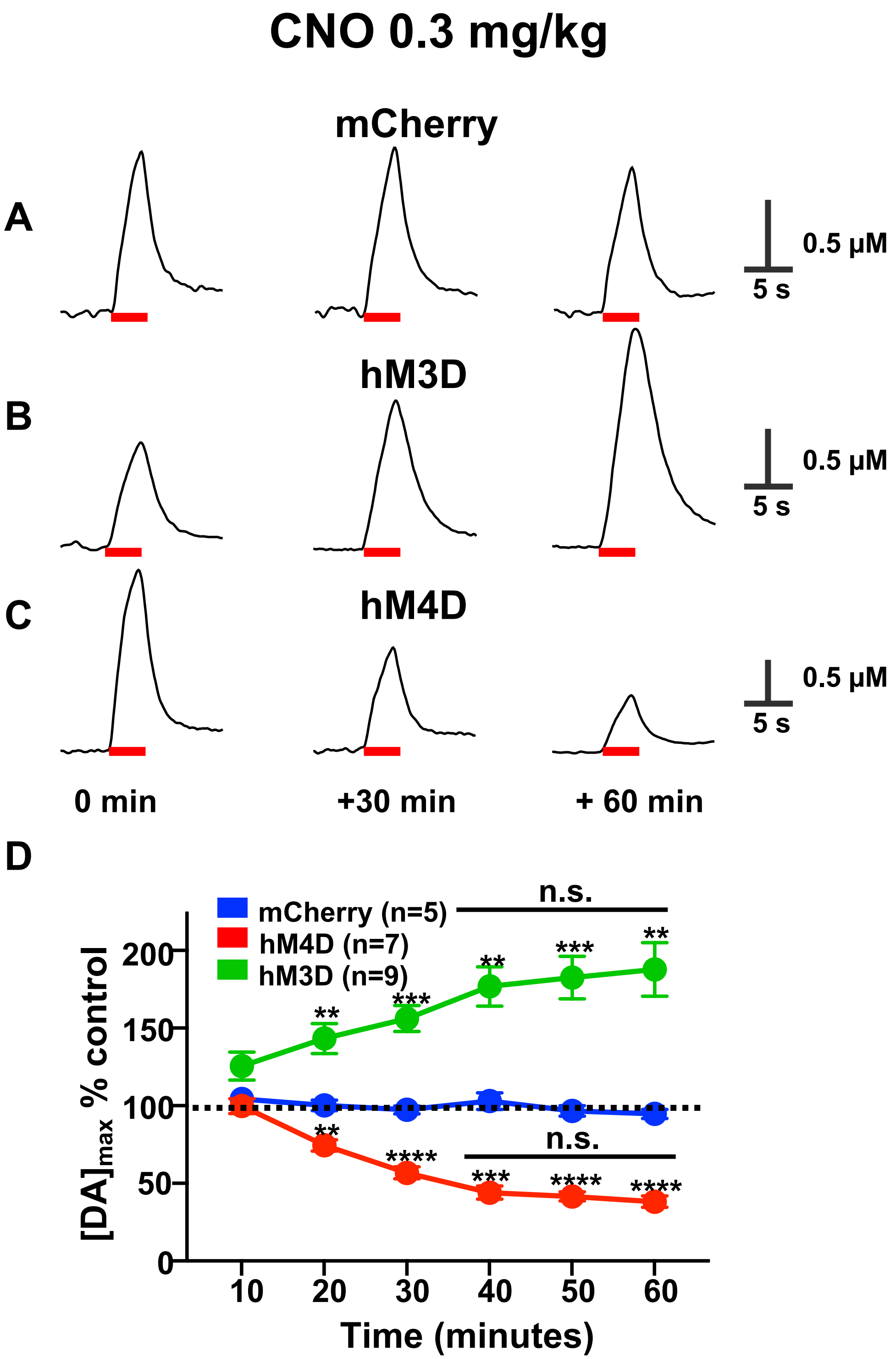


**Supplementary Figure 2:** Representative % concentration changes of $[DA]_{max}$ vs. time traces recorded in the OT evoked by electrical stimulation (20 Hz, 60 pulses) of the VTA following systemic administration of 0.3 mg/kg CNO in mCherry (**A**), hM3D (**B**), and hM4D (**C**) rats before as well as 30 and 60 minutes after CNO. (**D**) average changes in electrically evoked [DA]max recorded every 10 min after CNO administration. Electrical stimulation is denoted by red bar. n.s., not significant, **p < 0.01, ***p < 0.001, ****p < 0.0001.

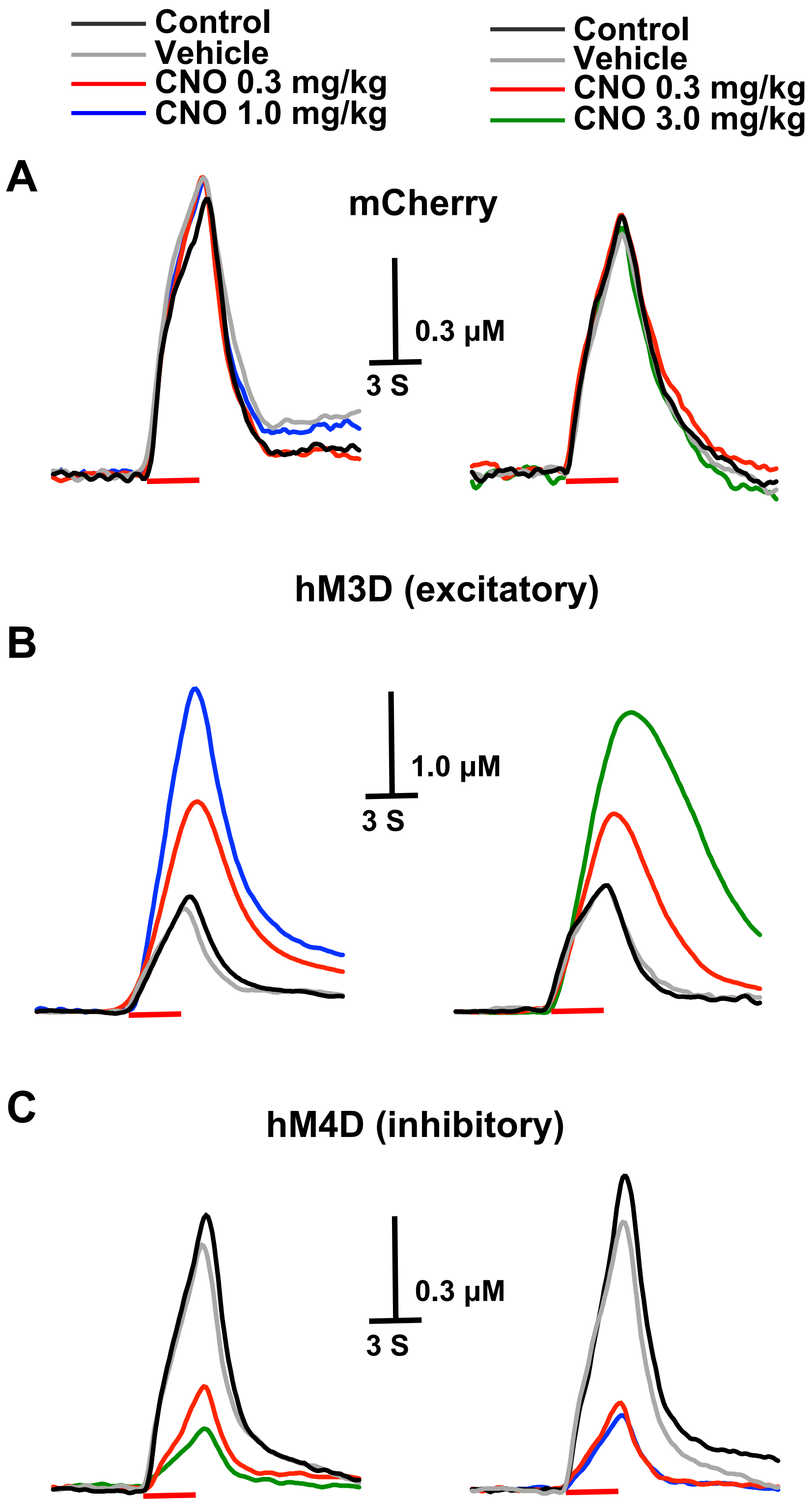


**Supplementary Figure 3:** Representative concentration vs. time traces for DA in the OT evoked by electrical stimulation of the VTA (20 Hz, 60 pulses, red line) following systemic administration of vehicle followed by different doses of CNO (0.3 and 1.0 mg/kg or 0.3 and 3.0 mg/kg) in mCherry (**A**), excitatory hM3D (**B**), and inhibitory hM4D (**C**) rats.
